# No evidence for a link between childhood (6-10y) cellular aging and brain morphology (12y) in a preregistered longitudinal study

**DOI:** 10.1101/2023.08.18.553696

**Authors:** Evie Henrike Brinkman, Roseriet Beijers, Anna Tyborowska, Karin Roelofs, Simone Kühn, Rogier Kievit, Carolina de Weerth

**Affiliations:** Donders Institute for Brain Cognition and Behaviour, Centre of Cognitive Neuroimaging & Radboudumc; Behavioural Science Institute, Radboud University, Donders Institute for Brain Cognition and Behaviour, Centre of Cognitive Neuroimaging & Radboudumc; Behavioural Science Institute, Radboud University, Nijmegen & Donders Institute for Brain Cognition and Behaviour, Centre of Cognitive Neuroimaging; Behavioural Science Institute, Radboud University, Donders Institute for Brain Cognition and Behaviour, Centre of Cognitive Neuroimaging; Lise Meitner Group for Environmental Neuroscience, Max Planck Institute for Human Development, Berlin, Germany & University Medical Center Hamburg-Eppendorf, Germany

**Keywords:** cellular aging, telomere length, epigenetic age, brain structure, paediatric imaging

## Abstract

Animal studies show that early life environmental factors, such as stress and trauma, can have a significant impact on a variety of bodily processes, including cellular aging and brain development. However, whether cellular wear-and-tear effects are also associated with individual differences in brain structures in humans, remains unknown. In this pre-registered study in a community sample of children (N=94, Mean age=12.71 years), we prospectively investigated the predictive value of two markers of cellular aging in childhood (at age 6 and 10) for brain morphology in early adolescence (age 12). More specifically, we associated buccal cell telomere length and epigenetic age in childhood to individual differences in adolescent whole-brain grey matter volume (GMV) including volumes of three regions of interest that have been found to be sensitive to effects of early life stress (i.e. amygdala, hippocampus, (pre)frontal cortex -PFC). Multiple regression analyses revealed no significant associations between childhood cellular aging (at 6 and 10 years) and early adolescent brain morphology. Exploratory Bayesian analyses indicated moderate to strong evidence for the null-findings. These results suggest that although our sample is modest, the associations between middle childhood cellular aging and early adolescent brain morphology are, if they do exist, likely not particularly large in community children. Future work should investigate whether these effects are similarly absent in large samples, in samples with a higher risk profile and in samples characterized by different age ranges.

**Highlights (3-5):**

- Investigation of cellular aging in relation to brain morphology in a community sample (N=95)
- Epigenetic aging and telomere shortening were not associated with brain structure
- Exploratory Bayesian Analyses reveal moderate to strong evidence for null findings
- No association was found between cellular aging and white matter volume

## 1. Introduction

Experiences early in life can shape the development of the body and the brain. Environmental factors, such as stress, toxins, and poor nutrition, can impact the wear-and-tear in cells as reflected by changes in markers of cellular aging. Notably, many studies have suggested similar associations between environmental risks and differences in brain morphology, suggesting that the wear and tear effects may ultimately manifest at the level of brain structure too (Colich et al., 2020). The goal of the current pre-registered study is to investigate whether markers of cellular aging (i.e. telomere length and epigenetic age) in childhood can predict brain morphology in early adolescence. Individual differences in adolescent brain morphology have been shown to predict later phenotypic outcomes such as aggression, emotion regulation, and episodic memory (see meta-analysis by Colich et al., 2020; or Dufford et al., 2019; el Marroun et al., 2016; Ghetti & Bunge, 2012; Hanson et al., 2010; Tyborowska et al., 2018). Given their roles in stress-related symptomatology, we will focus on the amygdala, hippocampus, and the PFC as specific regions of interest, in addition to whole brain morphology.

One marker of cellular aging is telomere length (Darrow et al., 2016; Harley et al., 1992). At the end of each chromosome is the telomere, a cap which protects the chromosome’s ends from degradation (Stewart et al., 2012).With each cell division the telomere shortens, resulting in telomere attrition with cellular age. Over time and with chronological aging, telomeres are steadily shortened, varying in rate throughout the lifespan (Vaiserman & Krasnienkov, 2021). Therefore, the length of the telomeres can be seen as a biological marker of cellular aging, with shorter telomere lengths reflecting older ages (Turner et al., 2019). Eventually, the telomeres will become too short and the cell will go into reproductive senescence (Sikora et al., 2021; Stewart et al., 2012). Importantly, telomeres in senescent state show a characteristic secretion pattern known as the senescence-associated secretory phenotype (SASP), including many cytokines and growth factors (Sikora et al., 2021). The secretion of these cytokines leads to an inflammatory state in the body, which has been extensively positively related to brain development as well as to accelerated aging (Sikora et al., 2021)

Indeed, first evidence from a cross-sectional study in 389 children (age 6-14) suggests that telomere lengths are associated with spontaneous activity in two main hubs of the default mode network (DMN): the posterior cingulate cortex (PCC) and the medial prefrontal cortex (mPFC) (Rebello et al., 2019). Moreover, previous research has linked activity in the DMN and telomere lengths to internalizing and externalizing problems (Davis et al., 2022; Farina et al., 2018; Sato et al., 2015). Shorter telomeres at age 6 were found to predict more self-reported internalizing and externalizing problems at age 10 in the sample of 193 community children from the current study (Beijers & Daehn, et al., 2020). These finding suggest an association between telomere length and properties of brain structure and function, but studies investigating this link in humans are scarce.

Another marker of cellular aging is epigenetic age. Epigenetic processes can be seen as the interplay between an individual’s environment and molecular biology (Hoare et al., 2020), and refer to the regulation of genome activities and gene expression by molecular modifications on the DNA. Literature shows that aging has an effect on the genome-wide DNA regulations, especially on DNA methylation levels. Therefore, the pattern of DNA methylation can estimate the age of the DNA source, not only reflecting the chronological age but also the biological age. This biological age can be captured in the form of an ‘epigenetic’ clock, a tool used to determine biological (i.e. epigenetic) age through determination of the methylation patterns of the DNA (Horvath, 2013; McEwen et al., 2020). To calculate the epigenetic clock in paediatric samples, the Paediatric Buccal cell Epigenetic (PedBE) clock is used, as it is the most accurate in predicting epigenetic age in children (McEwen et al., 2020).

Research has suggested that epigenetic processes, especially the methylation of DNA, play a mechanistic role in neurodevelopment and cell differentiation (Moore et al., 2013; Unnikrishnan et al., 2019). Additionally, there is growing evidence for the impact of early life events on methylation (Hoare et al., 2020; Horvath & Raj, 2018; Marini et al., 2020). For examples, the large study on 973 adults by Marini et al. (2020) showed that early life adversity may alter these normal methylation processes, leading to accelerated aging of cells. Consequently, this alteration of methylation patterns can lead to a deviation of the biological age from the chronological age, also known as epigenetic age acceleration (EAA) (Horvath, 2013; McEwen et al., 2020)

There is a dearth of studies on the association between epigenetic age acceleration and brain morphology. One recent study in young adolescents (N=44) from low income households found accelerated epigenetic age to be associated with alterations in brain morphology. More specifically, accelerated epigenetic age was associated with decreases in regional cortical thickness (Hoare et al., 2020). Another study in 4.5-year-old children (n=158), both with and without maltreatment experiences in the first 6 months of life, found that accelerated epigenetic aging was related to internalizing disorders and exposure to maltreatment (Dammering et al., 2021). Moreover, a study with a sample of 193 community children from the current study, epigenetic age acceleration was shown to be associated with internalizing behaviour, such that internalizing behaviour in 2.5-year old children predicted EAA at age 6 years, which in turn predicted internalizing behaviour at age 10 years (Tollenaar et al., 2021a). Both Dammering et al. (2021) and Hoare et al. (2020) argue that the stress response could underlie the association between accelerated epigenetic aging and brain morphology and functioning. They hypothesized that adversity early in life triggers a dysregulation of the stress system, which leads to dysregulated production of the stress hormone cortisol. Dammering et al. (2021) showed that epigenetic age acceleration was likely caused by such glucocorticoid dysregulation, as the CpG sites (DNA sequences where methyl groups bind) used for the PedBE clock are highly sensitive to glucocorticoids, which implies that glucocorticoids have an impact on the methylation of the DNA. This stress regulation of the epigenetic age would then cause a more rapid maturation of the brain (Dammering et al. (2021)).

In this study we will prospectively assess the predictive value of telomere length and epigenetic age acceleration in childhood (age 6 and 10) on grey matter brain volume (GMV) in early adolescence (age 12). At age 12, individual differences in GMV development are likely to be particularly pronounced, as only some children will have reached their peak GM volumes (Bethlehem et al., 2021). Brain regions that continue to develop into middle childhood, namely the amygdala, hippocampus, and prefrontal cortex, are especially vulnerable to effects of early life negative experiences (Romeo, 2017; Tyborowska et al., 2018). Indeed, both cross sectional as well as longitudinal studies have associated early life stress to reductions of these regional GM volumes (Romeo, 2017; Tyborowska et al., 2018).

In sum, while there is accumulating evidence linking adverse early life events to both cellular aging and brain morphology (Colich et al., 2020; Rebello et al., 2019), it remains unclear whether signs of cellular aging can also be linked to brain morphology. Therefore, the aim of the current preregistered study is to investigate potential associations between two biomarkers of cellular aging (i.e. telomere length and epigenetic age), and brain structure at age 12. Specifically, we will look at telomere length and epigenetic aging at ages 6 and 10, which reflect the cellular aging differences between individual, as well as at how they change over time, from age 6 to 10, reflecting increased or decreased cellular aging within individuals. Brain structure will be determined in terms of whole-brain GMV (using SPM), as well as a closer investigation of three regions of interest hypothesized to be especially relevant as they have been associated to early life stress (i.e. amygdala, hippocampus, and (pre)frontal cortex -PFC). We hypothesized that shorter telomere lengths and higher epigenetic age will be associated with smaller GMV, on the whole-brain level and particularly for the amygdala, hippocampus, and PFC. Additionally, the analyses mentioned above will be done with white matter volume and different brain sub-regions as outcomes measures. Because of a lack of previous literature, these analyses are exploratory in nature and therefore no specific directional hypotheses have been formulated.

## 2 Materials and Methods

### 2.1 Participants and procedure

Following recommendations about research practices (Nosek et al., 2018; Wagenmakers et al., 2012), this study was preregistered at AsPredicted: https://aspredicted.org/blind.php?x=kv8cb9 (632397). This study used data from an ongoing longitudinal project, the BIBO study (Basale Invloeden op de Baby Ontwikkeling; Dutch for Basal Influences on Infant Development) project, which aims to investigate the influence of early environmental factors and individual characteristics on child development. The BIBO study originally comprised 193 healthy, community, mother-child dyads, that have been followed since pregnancy (see (Beijers et al., 2011), for information about the original study). Since 80.9% of the mothers attended college or university we consider the sample low-risk. When children were 6 and 10 years old, buccal swabs were collected by researchers, to obtain genetic material (telomere length and DNA methylation) (Asok et al., 2013; Beijers & Daehn, et al., 2020; Beijers & Hartman, et al., 2020; Drury et al., 2014; McEwen et al., 2020). For the 12-year BIBO collection wave, 159 children were still participating in the study and were invited for an fMRI scan. Children with braces were excluded from participation (N=30), and several children did not participate for other reasons (e.g., too busy, no interest; N=31), resulting in a group of 97 children taking part in the visit. Markers of cellular aging did not differ between the 23% of the children that did not participate and the 77% that did (see Table 1).

**Table 1.** Differences in cellular aging between children not participating in the MRI-scan and children participating in the MRI-scan.

|  | *Mean [SD]*<br>*(non-MRI*<br>*subsample; 23%)* | *Mean [SD]*<br>*(MRI*<br>*subsample;*<br>*77%)* | *t-value* | *p-value* |
| --- | --- | --- | --- | --- |
| *Telomeres at 6 years* | 1.12 [0.61] | 1.11 [0.53] | -.0122 | .903 |
| *Telomeres at 10 years* | 0.58 [0.33] | 0.62 [0.33] | -1.521 | .130 |
| *Epigenetic age at 6 years* | 6.72 [0.65] | 6.72 [0.67] | -.018 | .985 |
| *Epigenetic age at 10 years* | 11.15 [1.00] | 11.02 [1.64] | .033 | .947 |

A mock scanner was used to familiarize the children with the scanning environment before the MRI session. MRI data of 3 children were excluded from further analysis due to poor quality, resulting in a final study sample of 94 children, of which descriptive sample characteristics and study variables are summarized in Table 2. The children had an average chronological age of 12.71 (SD=0.3) at the MRI visit. The average chronological age at the time of buccal swab collection was 6.09 (SD= 0.24) years, and 10.09 (SD=0.29) years. This study was approved by the ethical committee of the Faculty of Social Sciences of the Radboud University and the local medical ethics committee (CMO region Arnhem – Nijmegen). The children participated voluntarily and gave oral assent, and their parents provided written informed consent prior to participation.

**Table 2.**
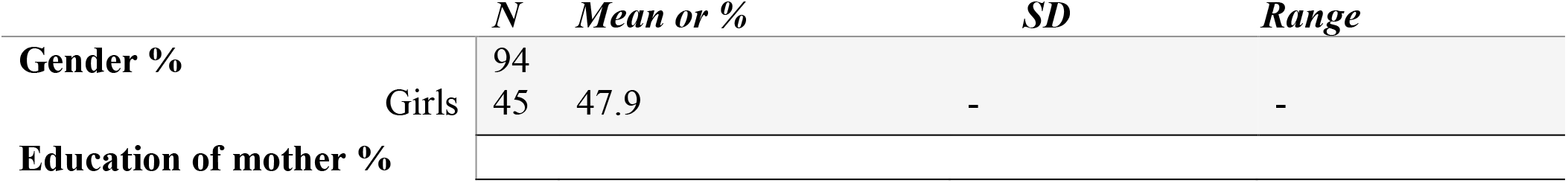
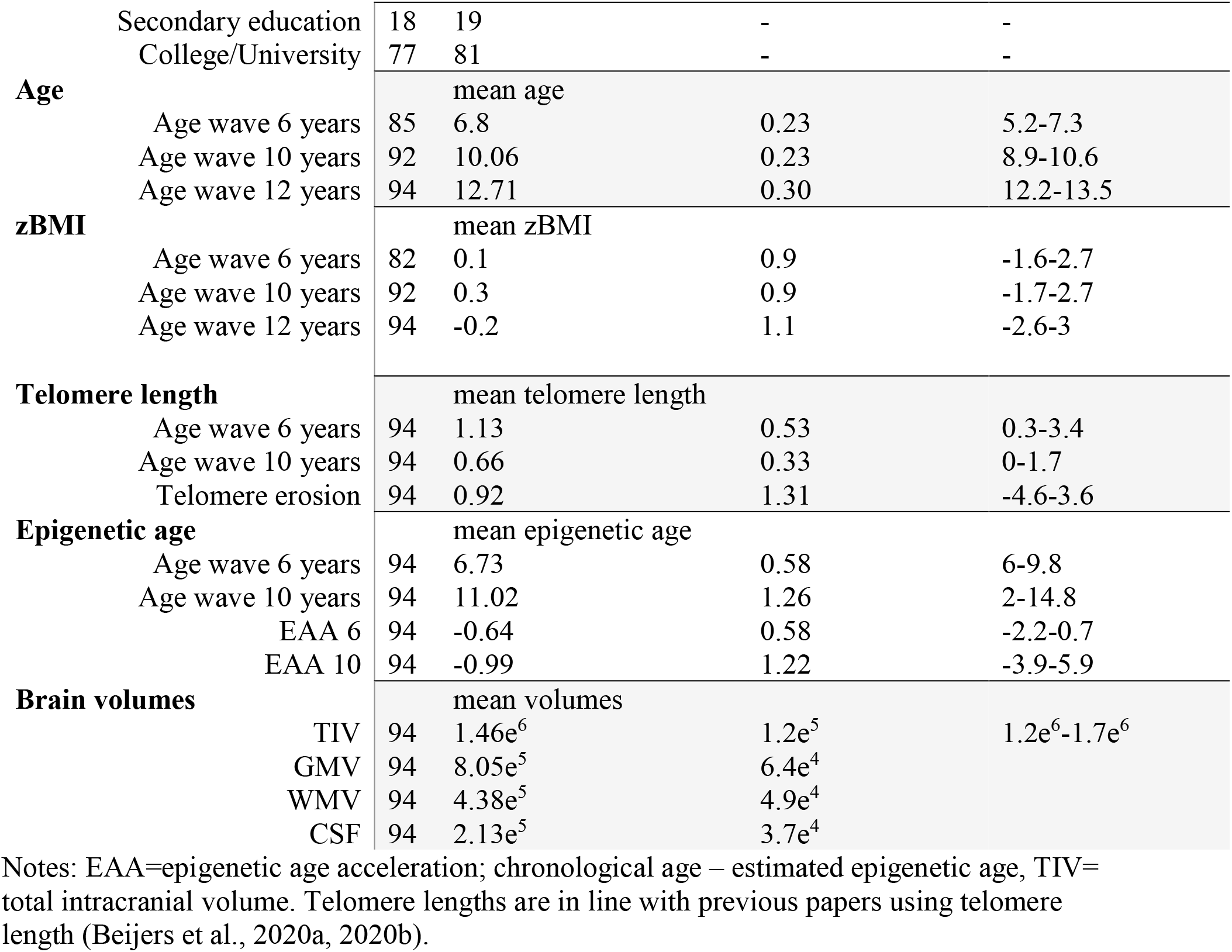
Overview of the children’s demographic characteristics and descriptive of the (raw) variables.

### 2.2 Measures

#### 2.2.1 Telomere length

DNA was extracted from buccal epithelial cells collected at age 6 (M=6.09, SD=0.24) and at age 10 (M=10.09, SD=0.29) using QIAamp DNA Mini Kit (Qiagen, Germany), and was quantified using Quant-iT PicoGreen reagent (Thermo Fisher Scientific, Qiagen). A quantitative PCR protocol was used to perform telomere length assays (see Beijers & Daehn et al., (2020a) and Beijers & Hartman et al., (2020b).

Telomere length is operationalized using the formula 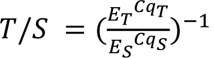, where E is the efficiency of exponential amplification for reactions targeting the telomere single-copy gene respectively, and Cq_T/S_ is the cycle at which a given replicate targeting telomeric content or the single-copy gene reaches the critical threshold of fluorescence quantification. The same threshold was used for all assays (36B4 and telomere). As samples were run in triplicate, the mean telomeric content 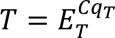 and mean genome copy number 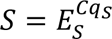 across replicates was used for calculating the T/S ratio. The mean was recalculated using two replicates when the replicates deviated from the mean telomeric content or mean genome copy number with more than 1.5% and was considered an outlier. Inter-assay variability was controlled for in line with Beijers & Daehn et al., (2020a) and Beijers & Hartman et al., (2020b).

Telomere lengths were corrected for age differences at the time of data collection by creating residuals. These were derived from regressing the telomere lengths of each assessment moment on the child’s age in months at that time point (Beijers & Daehn, et al., 2020; Beijers & Hartman, et al., 2020). Negative residuals indicate accelerated aging, as telomere lengths are shorter than expected, whereas positive residuals indicate slower aging, as telomere lengths are longer than expected (Fig. 1A-C).

**Figure 1.**
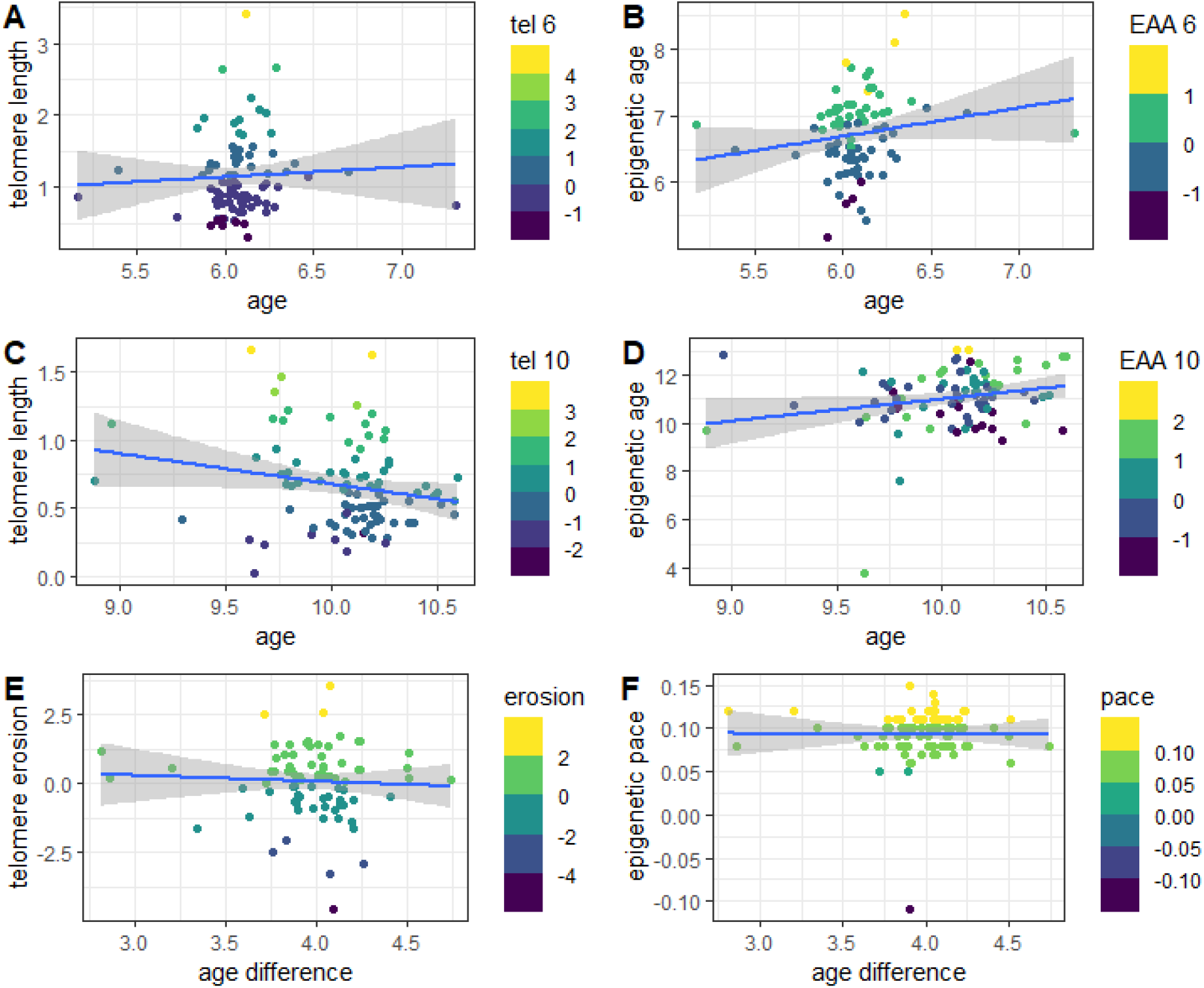
Distribution of telomere length and epigenetic age in the sample. **A**. The variance of residualized telomere length at age 6. **B**. The variance of epigenetic age acceleration at age 6 **C.** The variance of residualized telomere length at age 10. **D**. The variance of epigenetic age acceleration at age 10. **E.** The variance of telomere erosion between age 6 and 10 years. **F.** The variance of epigenetic aging pace between age 6 and 10 years. **A-D** The bluer colours indicate accelerated cellular aging, and the more yellow colours indicate decelerated cellular aging. **E+F** The more yellow colours indicate faster cellular aging pace, and the bluer colours indicate slower cellular aging pace. Note: The outliers did not affect further analyses

Furthermore, a measure of telomere erosion between age 6 and 10 years was created (see Beijers & Daehn et al., (2020a) and Beijers & Hartman et al., (2020b). Here, positive values reflect telomere erosion acceleration between age 6 and 10 (Fig.1E). Negative values reflect telomere erosion deceleration between age 6 and 10.

#### 2.2.2 Epigenetic age acceleration

DNA was extracted from buccal cells collected age 6 (M=6.09, SD=0.24) and at age 10 (M=10.09, SD=0.29) using QIAamp DNA Mini Kit (Qiagen, Germany). The Infinium MethylationEPIC array (Illumina, USA) was used for the description of the genome wide DNA methylation, necessary for the determination of the epigenetic age. Thereafter, the Minfi package in R was used for signal extraction, data quality, and pre-processing of the raw data. Epigenetic age was calculated using the newly developed Paediatric-Buccal-Epigenetic (PedBE) clock (McEwen et al., 2020). The PedBE clock is derived from DNA methylation at 94 CpGs, sharing 1CpG with the Horvath clock (McEwen et al., 2020).

For validation of the age estimation of the PedBE clock in our sample, a Pearson correlation was conducted between estimated epigenetic age and chronological age and found a correlation of r=.225 (p<0.05) at age 6 and a correlation of r=.210 (p<0.05) at age 10. In addition, epigenetic age at age 6 was significantly correlated with epigenetic age at age 10 (r=.568, p=<0.01), suggesting that our quantification of epigenetic age is relatively consistent. The epigenetic age acceleration at age 6 and age 10 were operationalized as the residuals from a linear model regressing PedBE-derived estimates of epigenetic age on chronological age in months at the moment of data collection. A positive value reflects higher than expected epigenetic age, thus epigenetic age acceleration (EAA), whereas a negative value reflects lower than expected epigenetic age, thus epigenetic age deceleration (Fig. 1B-D).(McEwen et al., 2020; Tollenaar et al., 2021b).

The pace of epigenetic age acceleration between age 6 and age 10 was operationalised as the difference in raw DNA methylation estimates over time (T2-T1) (Wolf et al., 2019). Values greater than 1.0 suggest an accelerated pace of epigenetic aging relative to the chronological aging, while values less than 1.0 suggest slower pace of epigenetic aging relative to chronological aging (Fig. 1F).

#### 2.2.3 Brain data

Brain MRI data was acquired using a 3T MAGNETOM PrismaFit MR scanner (Siemens AG, Healthcare Sector, Erlangen, Germany) with a 32 channel-coil. The children were scanned in the supine position. An MPRAGE sequence (TR = 2300 ms, TE = 3.03 ms, 192 sagittal slices, voxel size = 1.0 x 1.0 x 1.0 mm, FOV = 256 x 256 mm) was used to acquire whole brain T1-weighted images.

### 2.3 Data pre-processing

#### 2.3.1 Biological data pre-processing

We used the Markov Chain Monte Carlo procedure in SPSS to impute data of 9 children who were missing data from either the 6-year or 10-year measurement waves. The remaining participant was missing data of more than one predictor and was excluded from further analysis.

The six biological variables (telomere length at age 6 and 10, telomere erosion between age 6 and 10, epigenetic age at age 6 and 10, and epigenetic pace between age 6 and 10) were checked for violations of normality and outliers. Outliers were identified using the Multivariate Mahalanobis Distance (MD). In accordance with Tabachnick & Fiddell (2007), data of one participant was considered as an outlier, as it exceeded the critical chi-square value (degrees of freedom, df=6; the number of predictor variables in the model) at a critical alpha value of .001. This participant was excluded prior to further analysis.

Pearson’s correlations between the study variables and zBMI, and buccal cell count were evaluated to check the possibility of the latter two acting as confounding factors. No significant correlations were found between zBMI and the covariates of interest. Therefore, zBMI was not included as a covariate of no interest in further analyses. A significant correlation was found between buccal cell count at age 10 and epigenetic pace between age 6 and 10 (0.577, p=<.001, df= 92). Therefore, a residual of epigenetic pace was calculated using a regression of buccal cell count at age 10 on epigenetic pace.

#### 2.3.2 MRI data pre-processing

Raw structural T1-weighted images were checked for anatomical abnormalities, movement artefacts, and alignment to the anterior commissure. We performed the following pre-processing steps using Statistical Parametric Mapping 12 (SPM12), which is implemented in MATLAB (version 2019a). First, we used the Diffeomorphic Anatomical Registration Through Exponential Lie (DARTEL) algorithm to segment images into grey matter (GM), white matter (WM), and cerebrospinal fluid (CSF), and inter-subject registration of the GM images to a group average template image. Subsequently, GM images were normalized into Montreal Neurological Institute (MNI) space using a customized paediatric tissue probability map, which was created using the Template-O-Matic (TOM; version 1; http://141.35.69.218/wordpress/software/tom) toolbox. As a last step, grey matter images were smoothed using an 8×8×8 mm FWHM Gaussian kernel. Data quality after pre-processing was checked using the “Check Sample Homogeneity” function of the Computational Anatomy Toolbox 12 (CAT12), which indicated data of six children as potential outliers, of which data of two subjects was excluded from further analysis, after visual inspection. Total Intracranial Volume (TIV) was calculated as the sum of GM, WM, and CSF.

### 2.4 Statistical analyses

#### 2.4.1 Main analyses

To investigate the associations between accelerated cellular aging at age 6 and whole-brain GMV, a whole-brain multiple regression analysis was performed in SPM12 including the predictors at age 6, that is telomere length and epigenetic age. Additionally, a second multiple-regression analysis was performed including telomere erosion between 6 and 10 years, and epigenetic pace between 6 and 10 years, and whole-brain GMV at age 12 as outcome variable, to investigate the association between the longitudinal changes of the accelerated cellular aging markers and GMV. In both whole-brain multiple regressions, age, sex, and Total Intracranial Volume (TIV) were entered as confounders. To assess whole-brain statistical inference, the Threshold-Free Cluster Enhancement (TFCE) Toolbox in SPM12 was used to perform non-parametric permutation tests. For these permutation tests, a threshold of p<0.05 corrected for family wise error at a whole-brain level was used. The TFCE approach is especially advantageous for VBM data as it aims to enhance spatially contiguous signal without being dependent on threshold-based clustering. TFCE values at each voxel represented both spatially distributed cluster size and height information (Li et al., 2017; Smith & Nichols, 2009). Our ROIs (left and right amygdala, left and right hippocampus, and the PFC) were selected using the marsbar-AAL tool in SPM12 described by Tzourio-Mazoyer et al., (2002) The associations between accelerated cellular aging, both at age 6 as well as between age 6 and 10 years, and three ROIs were analyzed using multiple regression analyses in SPM12.

#### 2.4.2 Exploratory analyses

To exploratorily investigate the associations between accelerated cellular aging at age 10 and whole-brain GMV, a whole-brain multiple regression analysis was performed in SPM12 including the biomarkers of cellular aging at age 10, that is telomere length and epigenetic age.

Additionally, to exclude the possibility that the biomarkers of accelerated cellular aging are associated with brain regions that were not part of our a priori selection, all main analyses described above were repeated in an exploratory fashion with the following brain regions as outcome variables: the superior-, middle-, and inferior-gyrus of the frontal lobe, the precentral gyrus, the rectus, the inferior-, middle-, and superior-gyrus of the temporal lobe, the lingual gyrus, the fusiform gyrus, the insula, Heschl’s gyrus, the parahippocampal gyrus, the inferior- and superior-gyrus of the parietal lobe, the supramarginal gyrus, the postcentral gyrus, the precuneus, the inferior-, middle-, and superior-gyrus of the occipital lobe, the calcarine sulcus, the cuneus, the cingulum, the caudate, the putamen, the globus pallidus, and the thalamus. These brain regions were acquired using the SPM12 marsbar AAL-tool, as described by Tzourio-Mazoyer et al., (2002).

To explore whether accelerated cellular aging is linked to brain volume through a link with WMV, two multiple regression analyses were performed. The first included the predictors at age 6, that is telomere length and epigenetic age. The second analysis was performed including telomere erosion between 6 and 10 years, and epigenetic pace between 6 and 10 years, and WMV at age 12 years as the outcome variable, to investigate the association between the longitudinal changes of the accelerated cellular aging markers and WMV.

Lastly, for all multiple regressions Bayesian analyses were performed, to quantify the evidence in favor, or against, the regression models as compared to a null model. Results are expressed as a Bayes factor, which represents the relative likelihood of one model compared to another given the data and a prior expectation. This prior expectation was set as a default, noninformative JZS prior. The BayesFactor package from the open-source software package R was used to compute Bayes Factors (Morey & Rouder, 2015). For the interpretation of the evidential strength the description by Jeffreys (1961) was used, where a Bayes Factor <1/10 indicates strong evidence for the null-hypothesis, Bayes Factors >10 indicate strong evidence for H1, and a bayes factor of 0 indicates no evidence for either one (Jeffreys, 1961).

### 2.5 Deviations from the pre-registered study

In addition to the analyses described above, the pre-registration of this study describes the BrainAge, a model developed by Franke et al., (2010) that uses whole-brain neuroimaging data to reliably estimate the multidimensional aging pattern into one single value. The goal was to use BrainAge as a variable representing general aging of the brain. Therefore, a BrainAge model was created in R by regressing GMV (*M=8.05e^5^ mm^3^, SD=6.4e^4^ mm^3^*), white matter volume (*M=4.38e^5^ mm^3^, SD=4.9e^4^ mm^3^*), CSF (*M=2.13e^5^ mm^3^, SD=3.7e^4^ mm^3^*), and total intracranial volume (*M=1.46e^6^ mm^3^, SD=1.2e^5^ mm^3^*) onto chronological age. The BrainAge per individual was operationalised as the residual of this regression, in which positive values indicate older brains relative to the model-predicted age in this sample, whereas negative values indicate younger brains relative to the model-predicted age for an individual. As the variance in chronological age was relatively small in our sample, there was no correlation between GMV and age (.085). Therefore, the predicted BrainAge and GMV were significantly correlated (.482**). For this reason, BrainAge was excluded from further analyses.

## 3. Results

### 3.1 Descriptive analyses

Descriptive statistics are presented in Table 2 (untransformed data). Epigenetic age at age 6-years (M=7.76, SD= 0.69) and at age 10 (M= 12.44 SD=0.29) were significantly higher than the chronological age (t=-21.3 p=.001; t=-14.0 p=.001, respectively). Telomere length at 6 years of age (M= 1.11, SD= 0.56) and at 10 years of age (M= 0.62, SD=0.33) did not differ from each other (t=-.44, p=0.658), suggesting that between the ages 6 and 10 telomere length did not change in the same way for the group as a whole. Telomere length at age 6 and the pace of change were significantly correlated (r= -.664 p<0.01), which suggests that shorter lengths of the telomere at age 6 predict a higher pace of telomere erosion (see Supplemental Figure 3).

Table 3 shows the Pearson correlations between the study variables. Total Intracranial Volume (TIV) was significantly correlated with gender but insignificantly correlated with telomere length and epigenetic age. Interestingly, telomere length and epigenetic age were not correlated.

**Table 3.**
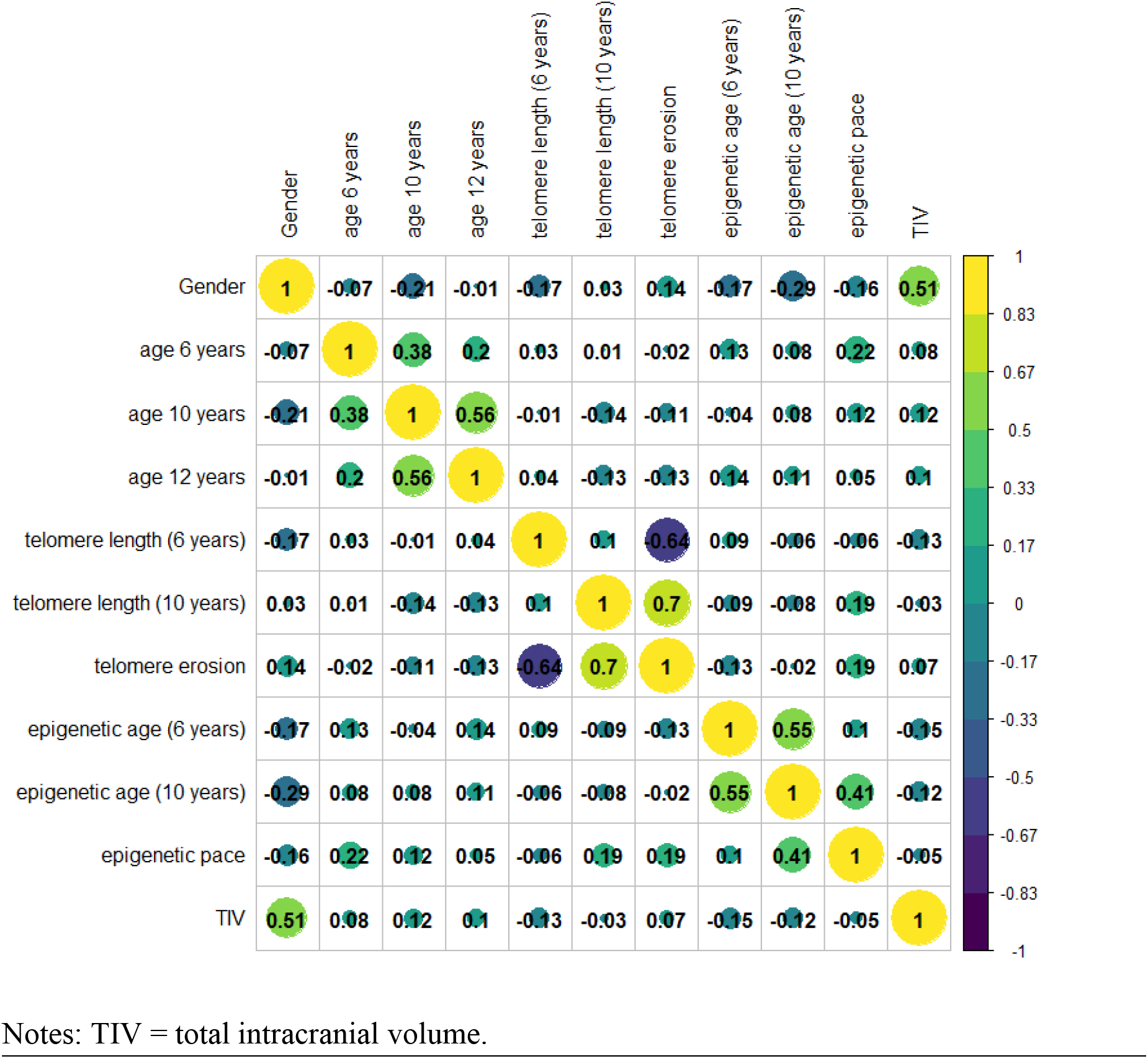
Correlogram representing the Pearson correlations between all study variables. Colors indicate different values of the correlation coefficient. The size of the circle is proportional to the correlation coefficients.

### 3.2 Main analyses

The analyses described under 3.2.1 to 3.3.2 were all carried out using the TFCE threshold of p<0.05 corrected for Family Wise error (FWE).

#### 3.2.1 Biomarkers of cellular aging at age 6 years

The whole-brain VBM analysis with telomere lengths and epigenetic age at 6 years did not yield any significant associations with whole-brain GMV (all corrected p-values above 0.998), nor with subregions of the brain thought to be associated with early life adversity, i.e., the amygdala, hippocampus, and PFC.

#### 3.2.2 Biomarkers of cellular aging at age 10 years

With respect to telomere lengths and epigenetic age at 10 years of age, the whole-brain VBM analysis did not yield any significant associations with these biomarkers and whole-brain GMV (all corrected p-values above 0.998), nor with subregions of the brain thought to be associated with early life adversity, i.e., the amygdala, hippocampus, and PFC.

#### 3.2.3 Changes in biomarkers of cellular age between 6 and 10 years of age

Regarding the changes in telomere length and epigenetic age between the age of 6 and 10, the whole-brain VBM analysis did not yield any significant associations with whole-brain GMV (all corrected p-values above 0.998), nor with the three subregions of interest (no suprathreshold values were found).

### 3.3 Exploratory analyses

#### 3.3.1 Exploratory brain regions

All the main analyses described above were repeated in an exploratory fashion with the brain regions described in the method section, acquired using the SPM12 marsbar AAL-tool (described by: Tzourio-Mazoyer et al. (2002)) as outcome variables. These analyses, using a threshold of p<0.05 corrected for Family Wise error at whole-brain level, also did not find significant associations.

#### 3.3.2 White matter volumes

Regarding both the telomere lengths and epigenetic age at 6 years of age, as well as the longitudinal changes in telomere length and epigenetic age between the age of 6 and 10, multiple regression analyses did not yield any significant associations with WMV (t=-1.312, p=0.193 and t=-1.148, p=0.254 respectively).

#### 3.3.3 Bayesian analyses

To be able to quantify whether the frequentist absence of effects was indeed evidence in favor of the null hypothesis, rather than an absence of precision or evidence either way, the main analyses were tested using Bayesian analyses. These analyses yielded Bayes Factors indicating a moderate to strong evidence for the null hypothesis (see Table 4). The full model with the predictors, telomere length and epigenetic age at age 6 was approximately 70 (bf=69.8) times less likely than the model including only the covariates of no interest (age, TIV, and gender), which suggests that evidence was found that indicates that it is unlikely that telomere length and epigenetic age predict GMV. Similarly, the full model with the predictors at age 10 was approximately 70 (bf=68.3) times less likely than the model including only the covariates. For both models including the predictors at age 6 or 10 years, the Bayes Factor indicated moderate evidence for the null hypothesis (bf= 0.172, and bf= 0.120 respectively). The model best predicting GMV, included only TIV (see Supplementary Material for all possible models).

**Table 4.** Bayes factor for selected models.

| **Telomere length and epigenetic age at age 6** |  | **Bayes factor** |
| --- | --- | --- |
| *Full model* |  | 5.888e+31 |
| Telomere length + epigenetic age |  | .172 |
| Telomere length |  | .361 |
| Epigenetic age |  | .373 |
| TIV, age, gender |  | 4.111e+33 |
| **Telomere length and epigenetic age at age 10** |  |  |
| *Full model* |  | 7.017e+31 |
| Telomere length + epigenetic age |  | .070 |
| Telomere length |  | .217 |
| Epigenetic age |  | .217 |
| TIV, age, gender |  | 4.111e+33 |
| **Telomere erosion and epigenetic pace between 6 and 10** |  |  |
| *Full model* |  | 6.0239e+31 |
| Telomere erosion + epigenetic pace |  | .120 |
| Telomere erosion |  | .284 |
| Epigenetic pace |  | .270 |
| TIV, age, gender |  | 4.111e+33 |

Notes: Full model= GMV~Telomere length+epigenetic age+TIV+ age+gender. GMV= grey matter volume, TIV= Intracranial Volume, age= age at 12 year measurement round.

## 4. Discussion

The current study investigated whether markers of cellular aging assessed in middle childhood, namely telomere length and epigenetic age acceleration, were associated with brain morphology in early adolescence in a low-risk sample. We hypothesized that shorter telomere length and higher epigenetic age acceleration at age 6 and 10 would be associated with smaller whole-brain grey matter, particularly in the three regions of interest (i.e. amygdala, hippocampus, and PFC) at age 12. Contrary to our expectations, we found no evidence for associations between the cellular aging markers and brain structure. These results were supported by exploratory Bayesian analyses, revealing Bayes Factors indicating moderate to strong evidence for the null findings. Finally, exploratory analyses inspecting associations between the two markers of cellular aging and both white matter as well as other subregions of the brain, not previously described in relation to the topic, delivered null findings as well.

One explanation for these findings points to the fact that the neural effects were tested two years after measuring cellular aging. Potentially, the associations between cellular aging and brain morphometry are only short-lived. The brain develops rapidly and is known to be vulnerable to environmental factors, particularly during its development in adolescence. It is possible that cellular aging processes at ages 6 and 10 did lead to short-term changes in the brain but that because of the brain’s plasticity at these ages, potential temporary impacts of cellular aging (f.i. through inflammation or glucocorticoid dysregulation) were reversed in the subsequent months or years. Such reversal mechanisms may particularly function in community samples in which the levels of stress, and hence the impact on cellular aging processes, are not as high as in high-risk samples where, in turn, effects on the brain may be less reversible and more cumulative in nature. To clarify these issues, longitudinal studies investigating both cellular aging as well as brain morphology over time are needed. Alternatively, it is possible that the effects of cellular aging processes on brain development may be more permanent, as they could possibly be considered programming effects. Accordingly, a possible alternative explanation for our null-results could be that the period from 6-10 years may be a period in which stress has less impact on the brain. Potentially, stress early in life may impact cellular aging which in turn affects brain developmental trajectories.

Another explanation for our null-results is related to sensitivity of group versus individual analyses; group-level analyses of alterations in brain structure may be less sensitive than analyses that account for individual brain development. Early developmental studies focused on deterministic models of brain development, assuming brain development proceeds via a prescribed blueprint that is innately specified in all individuals. Genome-wide association studies have shown that volumetric brain changes are heritable, and associated with substantial variability in brain volumes across different children of the same age (Brown, 2017). Besides, developmental studies show a link between puberty onset, ranging between the ages of 8 and 14, and brain maturation, such that higher pubertal developmental scores are associated with more mature brain development (Beck et al., 2023; Dehestani et al., 2023; Holm et al., 2023). Potentially, the (small) effects of cellular aging on the maturation of the brain are masked by the greater effect of inter-individual variation on brain maturation. However, due to the current study design, we cannot disentangle faster rates of maturation (different starting points, same endpoint) from discrete volumetric differences during early adolescence. Further longitudinal studies investigating the effects of cellular aging on individual brain development trajectories are therefore needed.

A final possible explanation for our null results could be that the associations might be very weak in a low-risk sample such as the one in this study, and we hence may not have had enough power to detect them with a sample of 94 adolescents. Indeed, Rebello et al. (2020) found only marginal relations between telomere length and brain connectivity in a study sample of 389 individuals, which did not survive strict Bonferonni corrections. A power analysis suggests that with a sample size of 94 children, an alpha of 0.05, and a power of 0.8, an effect size of f^2^ of 0.067 with a critical t= 1.66 can be found, which is in the moderate to large range for individual differences (Gignac & Szodorai, 2016). However, the results of the Bayesian analyses indicate a moderate to strong evidence for acknowledging the null-hypothesis given our uninformative prior. Together, the results of our study suggest that although our sample size is modest, the associations between middle childhood cellular aging and early adolescent brain morphology are, if they do exist, likely not particularly large in community children.

### 4.1 Strengths and limitations

This pre-registered study has several strengths, including the longitudinal design and measurement of two markers of cellular aging, namely telomere length and epigenetic age, of which epigenetic age was determined with the PedBE clock, a new model specifically developed for children (McEwen et al., 2020). T1 images were registered to a pediatric specific standard space, allowing for a more accurate registration. Using Bayesian analyses, we could symmetrically quantify evidence in favor of the absence of associations in our sample. However, some limitations should also be acknowledged. First, cellular aging was measured at ages 6 and 10, while brain morphology was measured at age 12 years. The lack of cellular aging measures at age 12 could be considered a limitation, as it was not possible to account for potential contemporary associations. A second potential limitation is that out of the potential pool of 128 children that were eligible to participate in the 12-year measurement, around 23% declined participation. However, because the cellular aging markers did not differ between participating and non-participating children, this does not appear to be a limitation that could have influenced the results. A final limitation is that while the markers for cellular aging were assessed at two childhood ages, the whole-brain GMV were only measured at 12 years, meaning we could not relate accelerated cellular aging to changes in brain structure over time.

## 5. Conclusion

In conclusion, we found no significant associations between childhood cellular aging (at 6 and 10 years) and adolescent brain morphology. Exploratory Bayesian analyses indicated moderate to strong evidence for the null-findings. These results point at the lack of a strong relation between markers of cellular aging and brain volume during childhood. Future studies might benefit from a longitudinal study design with cellular aging measures in early development (younger ages), biological and brain measures at the same age, as well as individual as opposed to group-level brain development trajectories.

## 6. Acknowledgements

CRediT roles:

- Conceptualization
- Data curation
- Formal analysis
- Funding acquisition
- Investigation
- Methodology
- Project administration
- Resources
- Software
- Supervision
- Validation
- Visualization
- Roles/Writing - original draft
- Writing - review & editing.
-

EB – Conceptualization, Formal analysis, Investigation, Roles/Writing – original draft

RB – Funding acquisition (BIBO), Data curation, Writing – review & editing, Data curation

AT – Methodology, MRI data acquisition, Data curation, Writing – review & editing

KR – Methodology, supervision MRI data acquisition, Writing – review & editing

SK – Methodology, Writing – review & editing

RK – Methodology, Investigation, Supervision, Roles/Writing – original draft

CW – Project administration, Conceptualization, Funding acquisition, Supervision, Investigation, Roles/Writing – original draft

## 7. Data + Code statement

The data of this study is part of an ongoing longitudinal study of which the data is still acquired and analyzed. Therefore, it cannot be made openly available in a public repository. Moreover, the parents of the participating children signed an informed consent that did not include the possibility of openly available data. However, for research purposes such as meta-analyses, it is possible to request the anonymized data by contacting: dr. Carolina de Weerth using a formal data sharing agreement where the goals of the project are outlined and the data transfer and potential co-authorships are described.

The analyses of this study were performed on software openly accessible, namely R (Download the RStudio IDE - RStudio) and SPM12 (SPM12 Software - Statistical Parametric Mapping (ucl.ac.uk)).

## 10. Supplementary Material

**Supplementary Figure 1.**
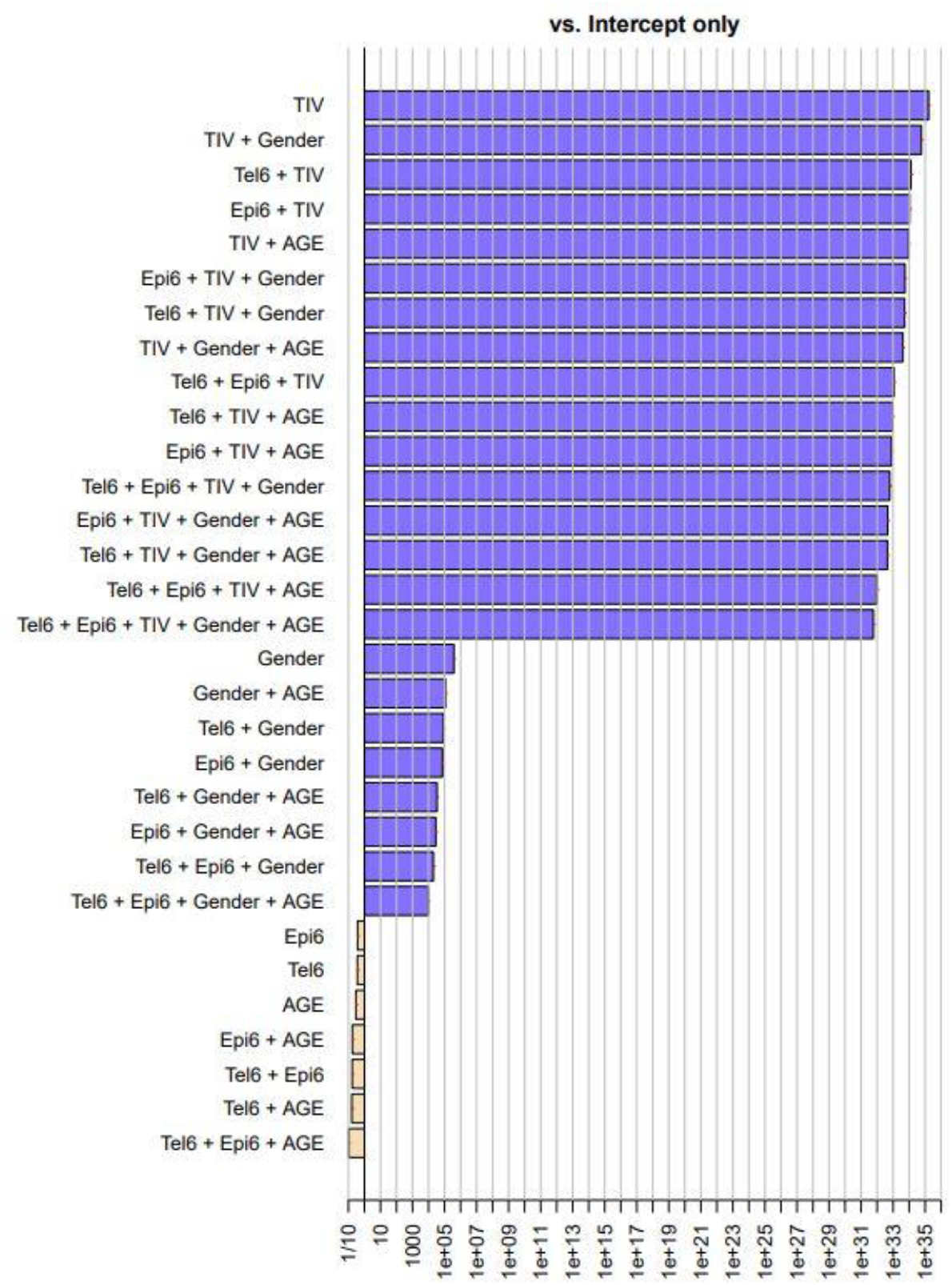
Bayesian analysis for all possible models predicting GMV at age 6 with the upper model presenting the model best predicting GMV and the lowest model presenting the worst model. Note: TIV= Total Intracranial Volume, AGE= age at MRI measurement round, Tel6 = telomere length at age 6, Epi6 = epigenetic age at age 6. Explain what the colours of the rows mean.

**Supplementary Figure 2.**
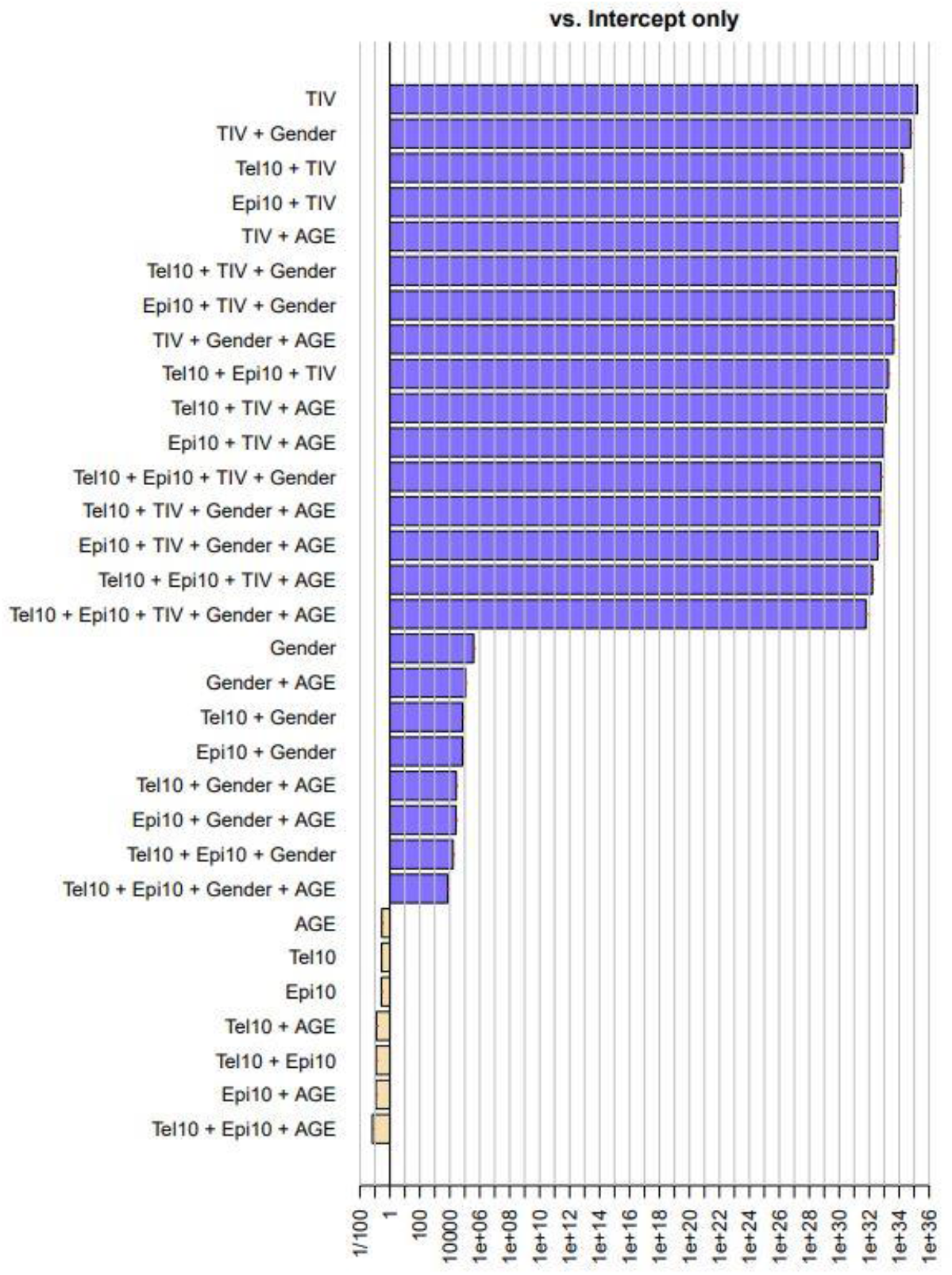
Bayesian analysis for all possible models predicting GMV at age 10 with the upper model presenting the model best predicting GMV and the lowest model presenting the worst model. Note: TIV= Total Intracranial Volume, AGE= age at MRI measurement round, Tel10 = telomere erosion between age 6 and 10, Epi10 = epigenetic pace between age 6 and 10. Explain colours.

**Supplementary figure 3.**
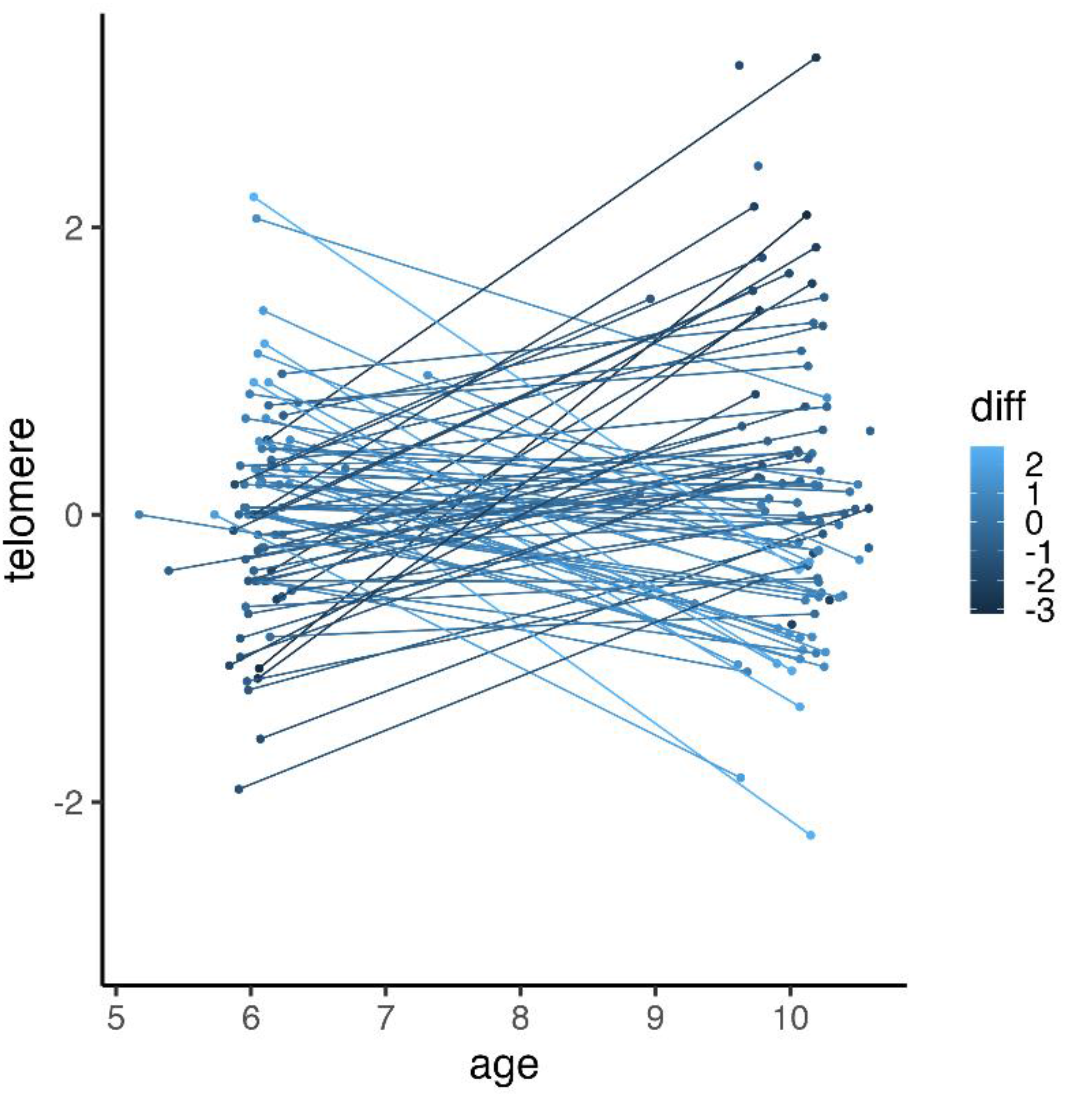
Difference in telomere length between age 6 and 10 years per individual. Individuals with shorter telomeres at age 6 than age 10 are depicted in dark blue, and individuals with longer telomeres at age 6 than age 10 are depicted in light blue.

