## Supplemental Figure 1 for "No evidence for a link between childhood (6-10y) cellular aging and brain morphology (12y) in a preregistered longitudinal study"

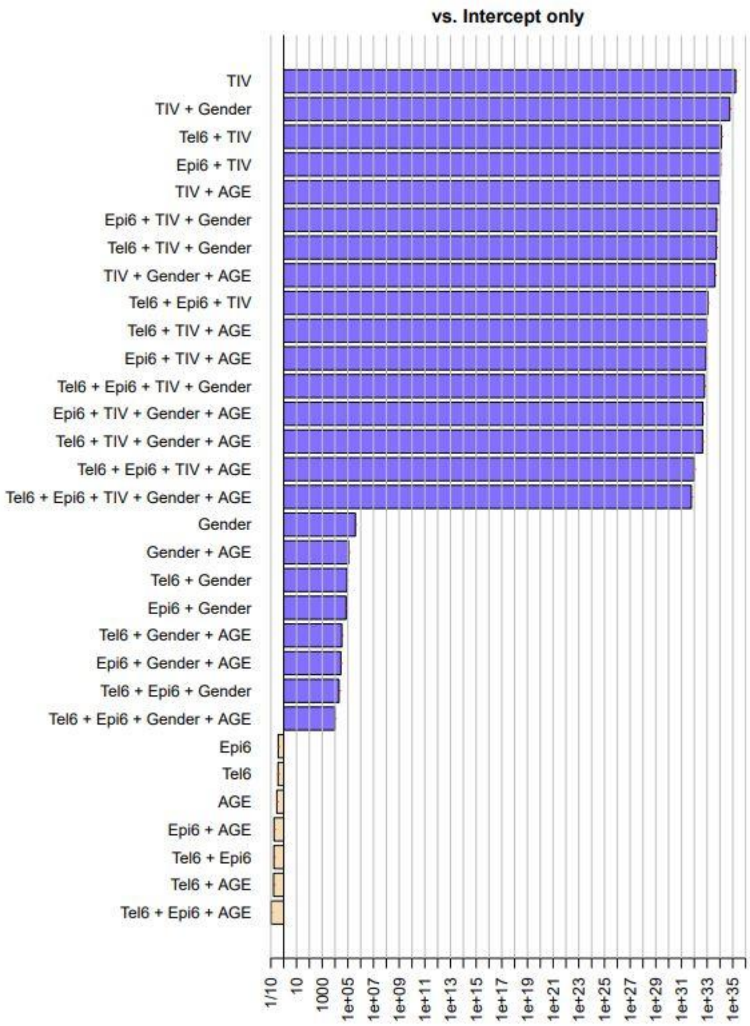


***Supplementary Figure 1.*** *Bayesian analysis for all possible models predicting GMV at age 6,* with the upper model presenting the model best predicting GMV and the lowest model presenting the worst model. Note: TIV= Total Intracranial Volume, AGE= age at MRI measurement round, Tel6 = telomere length at age 6, Epi6 = epigenetic age at age 6. Explain what the colours of the rows mean.
