## Supplemental Figure 2 for "No evidence for a link between childhood (6-10y) cellular aging and brain morphology (12y) in a preregistered longitudinal study"

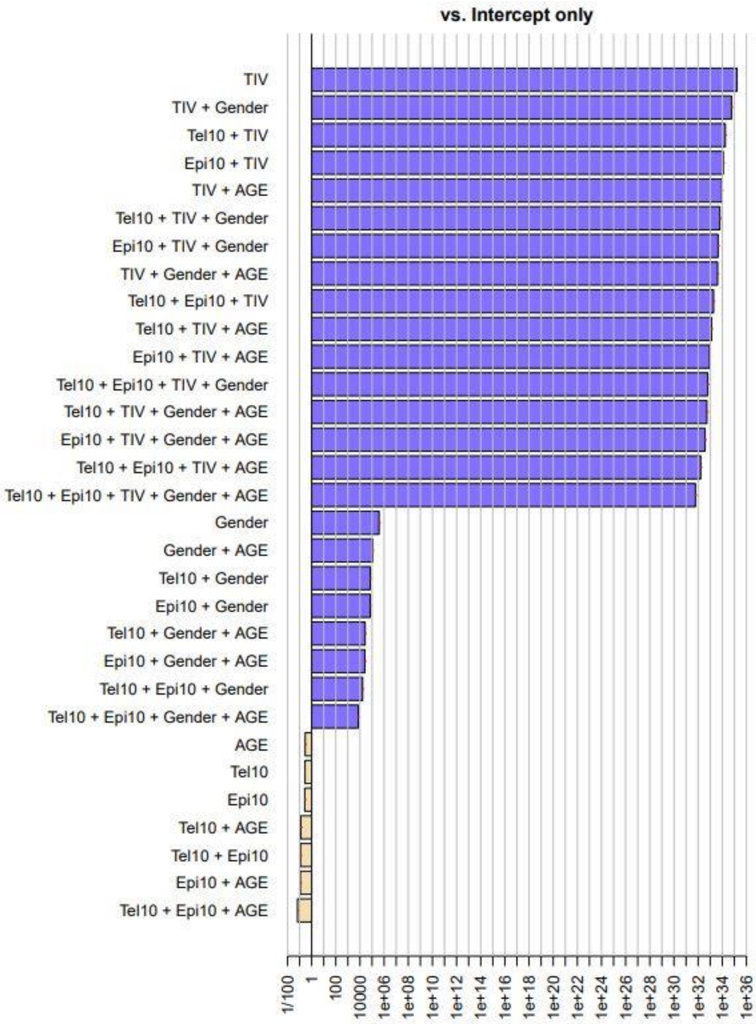


***Supplementary Figure 2.*** *Bayesian analysis for all possible models predicting GMV at age 10*, with the upper model presenting the model best predicting GMV and the lowest model presenting the worst model. Note: TIV= Total Intracranial Volume, AGE= age at MRI measurement round, Tel10 = telomere erosion between age 6 and 10, Epi10 = epigenetic pace between age 6 and 10. Explain colours.
