## Supplemental Figure 3 for "No evidence for a link between childhood (6-10y) cellular aging and brain morphology (12y) in a preregistered longitudinal study"

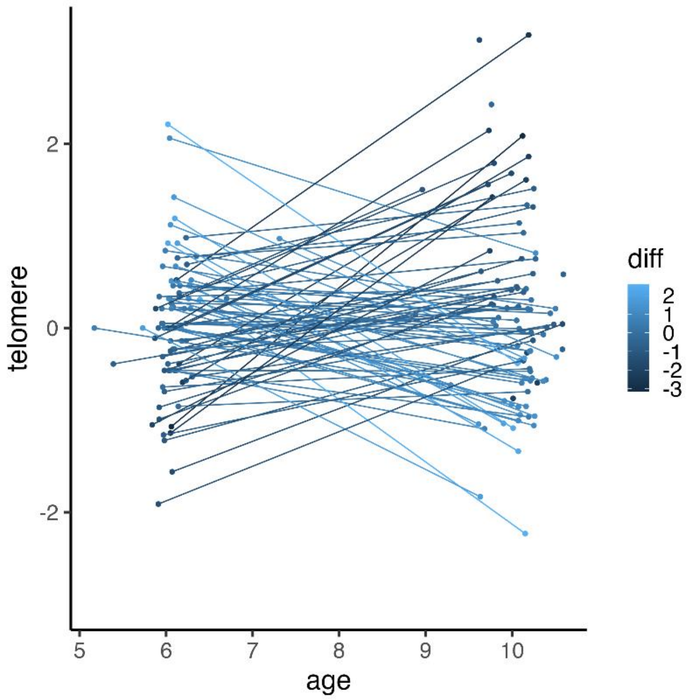


***Supplementary figure 3.*** *Difference in telomere length between age 6 and 10 years per individual.* Individuals with shorter telomeres at age 6 than age 10 are depicted in dark blue, and individuals with longer telomeres at age 6 than age 10 are depicted in light blue.
